# An interpretable integration model improving disease-free survival prediction for gastric cancer based on CT images and clinical parameters

**DOI:** 10.1101/2024.04.01.587508

**Authors:** Xiaoping Cen, Can Hu, Li Yuan, Xiangdong Cheng, Wei Dong, Run Zhou, Yuanmei Wang, Jiansheng Zou, Tianyu Lu, Huanming Yang, Yahan Tong

## Abstract

Preoperative prediction of disease-free survival of gastric cancer is significantly important in clinical practice. Existing studies showed the potentials of CT images in identifying predicting the disease-free survival of gastric cancer. However, no studies to date have combined deep features with radiomics features and clinical features. In this study, we proposed a model which embedded radiomics features and clinical features into deep learning model for improving the prediction performance. Our models showed a 3%-5% C-index improvement and 10% AUC improvement in predicting DFS and disease event. Interpretation analysis including T-SNE visualization and Grad-CAM visualization revealed that the model extract biologically meaning features, which are potentially useful in predicting disease trajectory and reveal tumor heterogeneity. The embedding of radiomics features and clinical features into deep learning model could guide the deep learning to learn biologically meaningful information and further improve the performance on the DFS prediction of gastric cancer. The proposed model would be extendable to related problems, at least in few-shot medical image learning.

**Key Points:**

- An integration model combining deep features, radiomics features and clinical parameters improved disease-free-survival prediction of gastric cancer by 3%-5% C-index.
- Embedding radiomics and clinical features into deep learning model through concatenation and loss design improved feature extraction ability of deep network.
- The model revealed disease progression trajectory and tumor heterogeneity.

## Introduction

Gastric cancer (GC) is the fifth most common malignant cancer worldwide and the fourth leading cause of global cancer deaths [1]. GC patients are usually diagnosed at an advanced stage with poor prognoses. The 5-year overall survival rate of GC is 30−40% after curative resection, whereas metastasis-related or relapse-related death remain a challenge for curative treatment [2]. Therefore, preoperative prediction of the disease-free survival (DFS) of gastric cancer is remarkably important for evaluating patient progression before treatment.

Efforts have been made on predicting the DFS of gastric cancer based on clinical reports and CT image-based radiomics. Discovered risk factors include age, tumor size, tumor stage, tumor location, and Lauren type. While some of these factors are determined postoperatively and are limited to precisely determine the disease progression of GC patients, some studies focused on the utility of preoperative CT images. Meanwhile, the development of radiomics enabled the extraction of rich information from images, showing considerable performance in many image-based prediction tasks. Radiomics features extracted from CT images suggested higher performance than clinical parameters in predicting DFS of gastric cancer, which can also predict chemotherapeutic benefits for stage II/III patients [3]. Multidetector-row computed tomography (MDCT)-based radiomics nomogram suggested that the integration of radiomics nomogram with clinical parameters improved the prediction performance in DFS prediction of GC [4]. Li et al. focused on stage II/III patients and developed a radiomics nomogram which integrated both intratumor and peritumor information for DFS and chemotherapy response prediction of GC [5]. However, as the author suggested, their model is limited to single-vendor CT scanners (GE).

Moreover, due to the high tumor heterogeneity, radiomics features are not enough to extract complex information from images. Deep learning has been widely applied in medical image analysis and cancer studies, showing considerable performance in gastric cancer prediction tasks [6, 7], also including a multi-task prediction of the DFS and peritoneal recurrence of GC on multi-institution data [8]. In order to further improve the performance, some studies have evaluated the performance of integrating deep learning features with radiomics features or clinical parameters [9-12]. However, linear regression models are usually applied, and the deep features are extracted offline, which limited the capability of deep learning-based feature extraction and lacks of flexibility and generalizability.

Some end-to-end integration methods were widely applied in multi-source data integration for medical image analysis. The common way is to train different networks for each source of data and then combine features from different sources at the last layer. In order to provide more intuitive integration, we proposed that the loss for each sub-networks would provide supervision on the main network, based on the principle of deeply supervised nets [13]. Therefore, we developed an interpretable framework for the integration of radiomics, clinical and deep features, and we hypothesized that radiomics features and clinical features as a kind of prior knowledge [14, 15], would provide supervision to the feature extraction, whereas deep learning could also correct the misinformation caused by the prior knowledge.

Similar methods were developed in previous studies. Wei et al proposed a strategy by first mapping radiomics features, clinical features into common feature space using variational autoencoder, and then integrate them with deep features, which showed improvement of performance in predicting the overall survival of hepatocellular carcinoma [16]. Zhou et al applied deeply supervised multi-modal integration scheme to predict the microvascular invasion using multi-phase contrast-enhanced MRI images [17]. However, the investigation on gastric cancer has not been performed.

In this study, we aim to evaluate the performance of integrating clinical parameters, deep learning features, and radiomics features in an end-to-end deep learning framework on gastric cancer DFS prediction. Besides, interpretation analyses are performed to reveal the biopathology mechanism revealed by the model.

## Material and Methods

### Dataset and Preprocessing

#### Study Population

The study obtained approval from the local institutional review board, and the requirement for patients’ informed consent was waived. All patients underwent surgical treatment within one month of contrast-enhanced abdominal CT examination. There were primarily 1001 patients diagnosed as GC. We applied the inclusion criteria followed to identify required patient samples: (1) postoperative pathologically confirmed gastric adenocarcinoma; (2) patients who underwent surgical treatment; (3) patients who underwent contrast-enhanced CT of the whole abdomen or the upper abdomen within one month before surgery; (3) no distant metastases prior to surgery. The exclusion criteria were as follows: (1) incomplete clinical or pathological information; (2) treatment was performed before surgery; (3) poor CT image quality or unrecognizable lesion; and (4) other concurrent malignancies. Finally, 214 patients were enrolled for this study.

#### CT image acquisition

All patients underwent contrast-enhanced abdominal CT using the multidetector row CT systems: BrightSpeed, Optima CT680 Series (GE Medical Systems); Siemens Somatom definition AS 64, Perspective (Siemens Medical Systems). The acquisition parameters were the following: detector configuration 128 × 0.6 mm; tube voltage, 120-130 kV; tube current, 150-300 mAs; reconstructed axial-section thickness 5 mm, slice interval 5 mm, pitch 0.6. The contrast agents were Ultravist (Bayer Schering Pharma, Berlin, Germany), Optiray (Liebel-Flarsheim Canada Inc., Kirkland, Quebec, Canada), and Iohexol (Beijing North Road Pharmaceutical Co. Ltd., Beijing, China). A contrast medium of 1.5 ml/kg was injected through the antecubital vein. The portal venous phase was performed at 50-60 s after injection of the contrast medium.

#### Tumor segmentation

The CT images of portal vein phase were exported in digital imaging and communications in medicine (DICOM) format from image storage and communication systems. Lesions were segmented using ITK-SNAP (version 3.8.0, http://www.itksnap.org). Two radiologists with over five years of experience in diagnosing abdominal diseases observed each CT image of the patient and performed delineation on the largest layer of the tumor. Here only 2D tumor region was delineated, as previous study suggested that 2D radiomic features revealed comparable performances with 3D features in characterizing GC [18], while one-slice 2D annotation saves time instead of whole-volume 3D annotation.

#### Radiomics feature extraction

A total of 944 radiomics features were extracted from the delineated ROI regions, including 2D shape, firstorder, glcm, glrlm, glszm, gldm, ngtdm features with original image or images after log and wavelet filtering [19]. Lasso cox regression model with 10-fold cross validation was applied to determine useful features for DFS prediction. The pyradiomics package [20] in Python 3.7.3 and glmnet package [21] in R were utilized for radiomics feature extraction and feature selection, respectively.

### The Proposed Method

#### The Overview of the Proposed Framework

The proposed framework was summarized in **Fig.1**.

**Figure 1.**
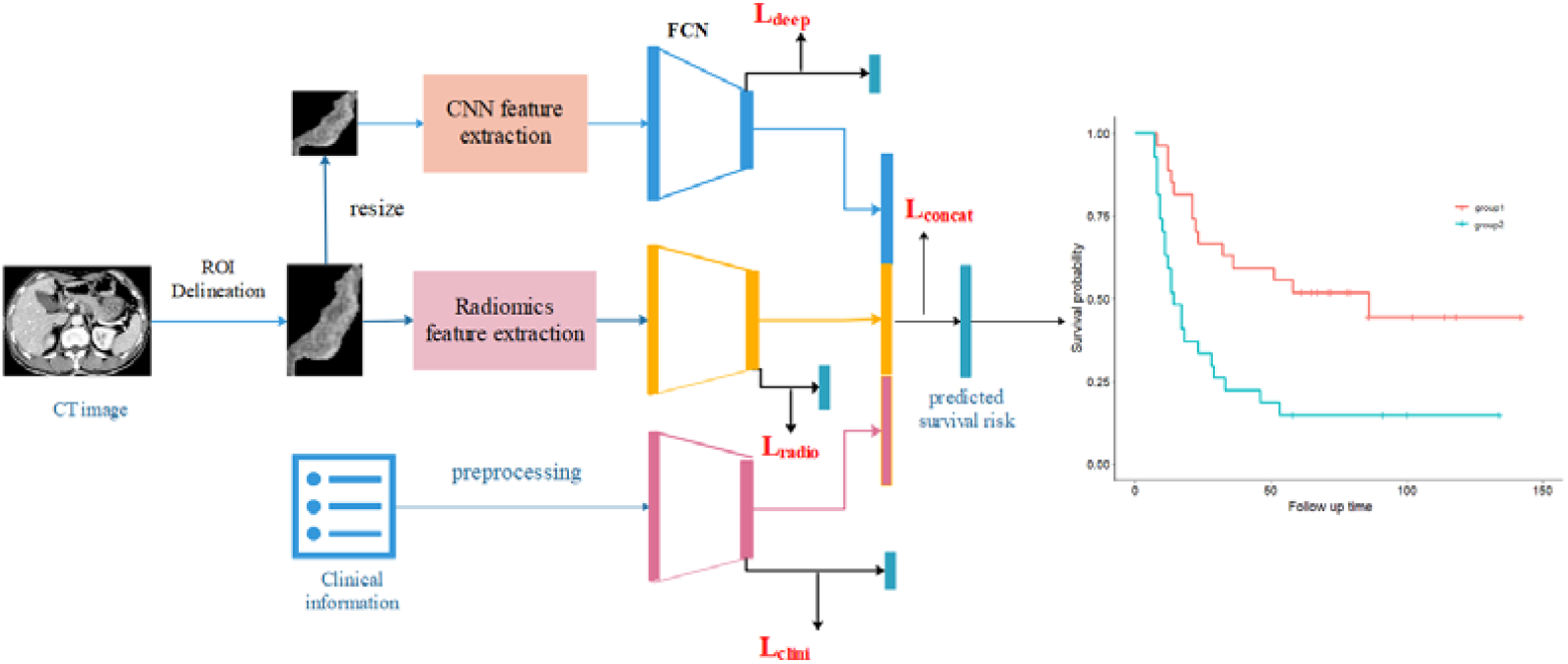
The proposed framework for integrating deep features, radiomics features and clinical features in a deep learning network

#### Descriptions and the Implementation

Images were resized into 28*28 as input of convolutional neural network (CNN). A ResNet50 model with pre-trained ImageNet weights for all convolutional layers was utilized to extract deep features. The ResNet50 model consists of 49 convolutional layers, followed by a fully connected layer. We added a fully connected layer into two layers to reduce the dimension of features into 256 dimensions, and the output 1000 dimension was changed into one dimension referring to the risk score predicted by the model. In order to embed the radiomics features and clinical parameters into the model, we first normalized the two sets of features, and then passed them into the deep network. A FCN layer was then utilized to embed the features into 250 and 6 dimensions, respectively. The three kinds of features, were then concatenated and formed 512 dimensions. Then, another FCN layer was then applied to predict risk score based on the features. We applied two kinds of strategies for calculating the loss value of the network: 1) simply using the loss value of the final 512 dimension features, named CON model; 2) combine the individual loss with the concatenated loss based on the design of deeply supervised nets [13], named DSN model. The DeepSurv [22] was used as loss function for cox regression. The DSN loss was defined as follows:

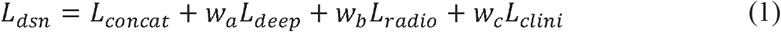

All experiments were implemented using the PyTorch machine learning library. The learning rate was set as 0.0001 and Adam optimizer was used to finetune all the model parameters. To alleviate the effect of overfitting problem to the model performance, we applied early stopping strategy to stop the iterations before the overfitting occurred [23].

### Interpretability Analysis

To assess whether the proposed models captured more information, we performed T-SNE analysis on both feature-level and sample-level using the Rtsne package [24, 25]. We utilized Grad-CAM visualization method [26] to visualize the area which the model has focused on.

### Statistical Analysis

The data set was randomly split into the training set (160 GCs) and validation set (54 GCs). Five-fold cross-validation was performed on the training set to determine the model parameters, and then the model was utilized for independent testing on the validation set. For each deep learning model, we performed five times training and testing to assess the stability of the models, and the final performances were calculated using mean ± sd. Once the predictions were made, Kaplan-Meier curve was plotted using the median value of the predicted risk score as cutoffs.

## Results

### Statistical description of clinical parameters

Table 1 presented the statistical description of clinical parameters among training set and validation set. Among the 214 GCs, there are 163 GCs obtaining relapse or metastasis after surgery. The DFS ranges from 7 to 177 months, with a medium value of 26 months.

**Table 1.**
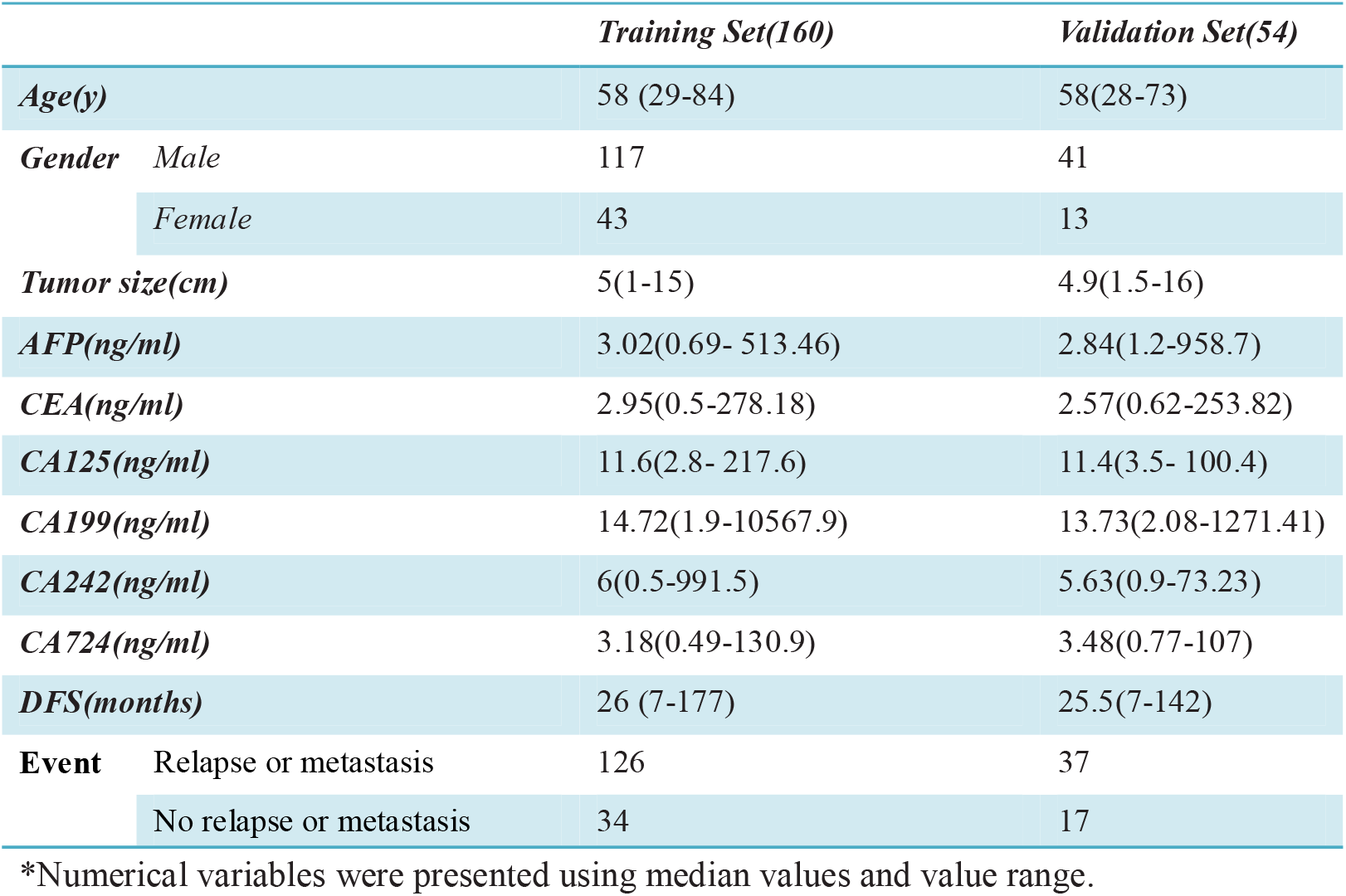
Statistical description of clinical parameters in training and validation set*.

### Model performance

We systematically compared the proposed framework with baseline models in Table 2. Radiomics model showed slightly better performance among the three baseline models. Lasso-based integration models did not improve the performance, while the proposed integration models showed improved C-index performance of 3%-5% in predicting DFS. Using the predicted score and the event label, we found that the DSN and CON model improved AUC of 10% in predicting event occurrence. However, while integrating multi-source information should lead to better performance, we found when integrating deep features, radiomics features with clinical features, the model did not obviously yield better performance than the model integrating only deep features and radiomics features. The result may be attributed to the fact that the clinical features are of low dimension, and that some measurements are imputed by mean values, which introduce significant noise into the model, overshadowing the valuable information it should capture.

**Table 2.**
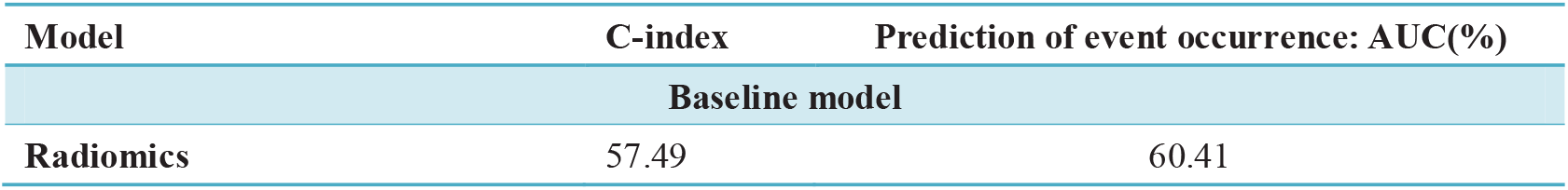
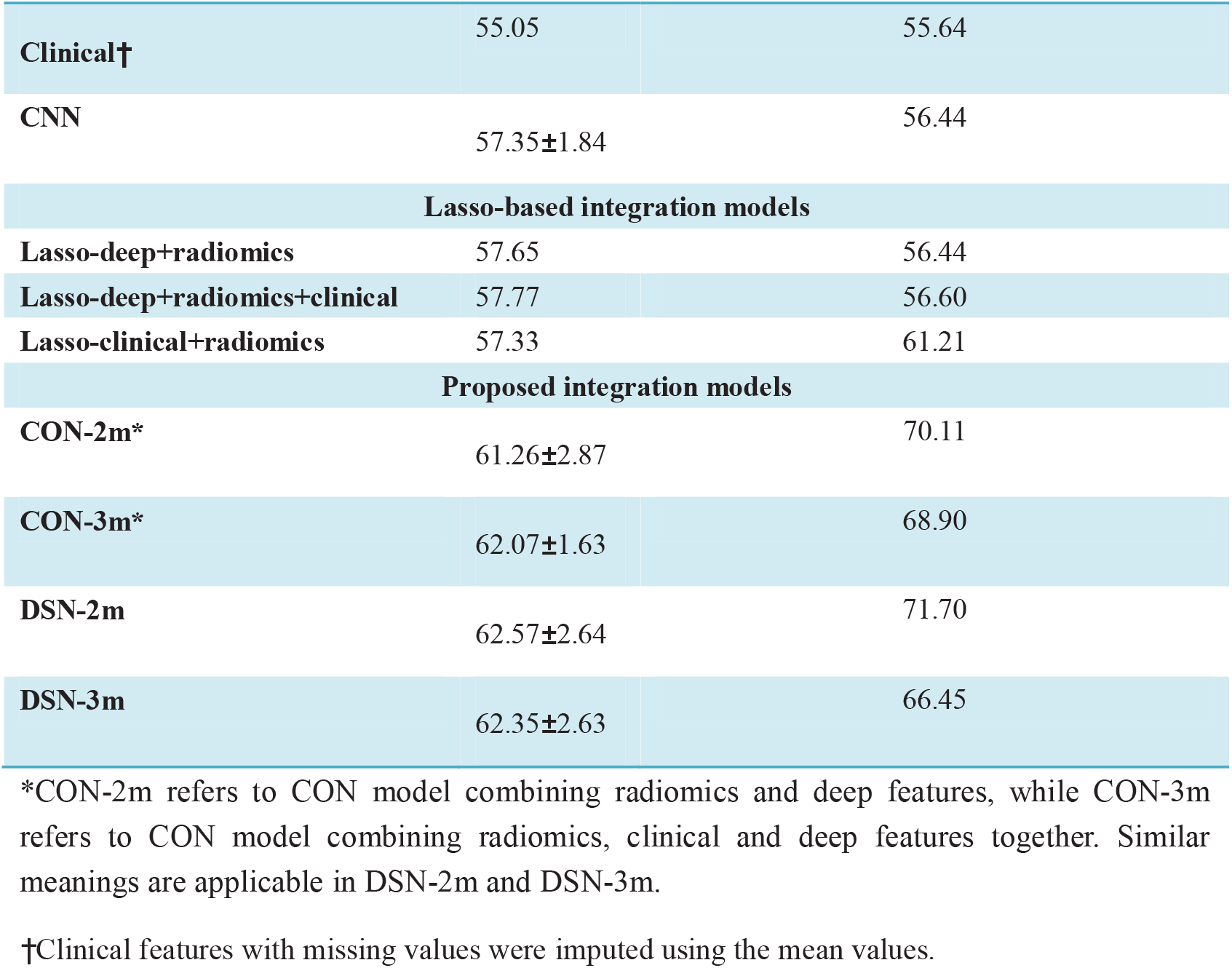
Performance comparison between different models.

In order to identify the time-dependent property of the models, we performed time-dependent ROC analysis for each model. According to Fig.2 and Table3, the proposed CON or DSN models present higher AUC among different time points for relapse prediction, and especially when the time is 21 months (1.75 year), the two models could perform significantly better than radiomics model. Besides, we found DSN model could predict patient relapse at eight-month period, which indicates DSN model could identify particularly aggressive gastric cancer that are not suitable for proposed treatment.

**Figure 2.**
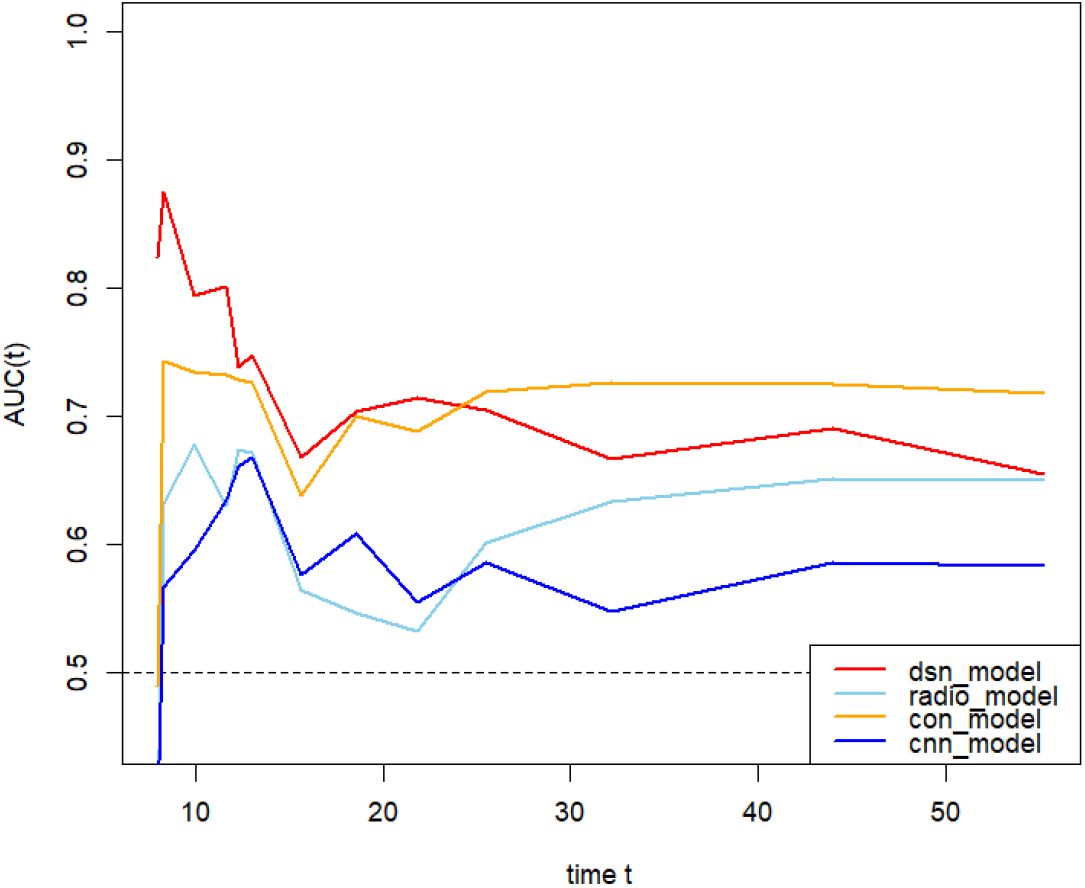
Time dependent ROC curves for different models

**Table 3.** Comparison between time dependent AUC for different models.

| Time(m) | radio vs dsn | radio vs con | cnn vs con | cnn vs dsn | con vs dsn |
| --- | --- | --- | --- | --- | --- |
| t=8 | <0.001 | 0.653 | 0.519 | <0.0001 | 0.019 |
| t=8.3 | 0.045 | 0.385 | 0.179 | 0.025 | 0.159 |
| t=21.85 | 0.026 | 0.040 | 0.127 | 0.057 | 0.689 |

Kaplan-Meier curves **(Fig.3)** also proved that the proposed DSN and CON models showed good differentiation ability for high-risk and low-risk patients, while the radiomics and CNN model showed no statistical significance in differentiating high-risk and low-risk patients.

**Figure 3.**
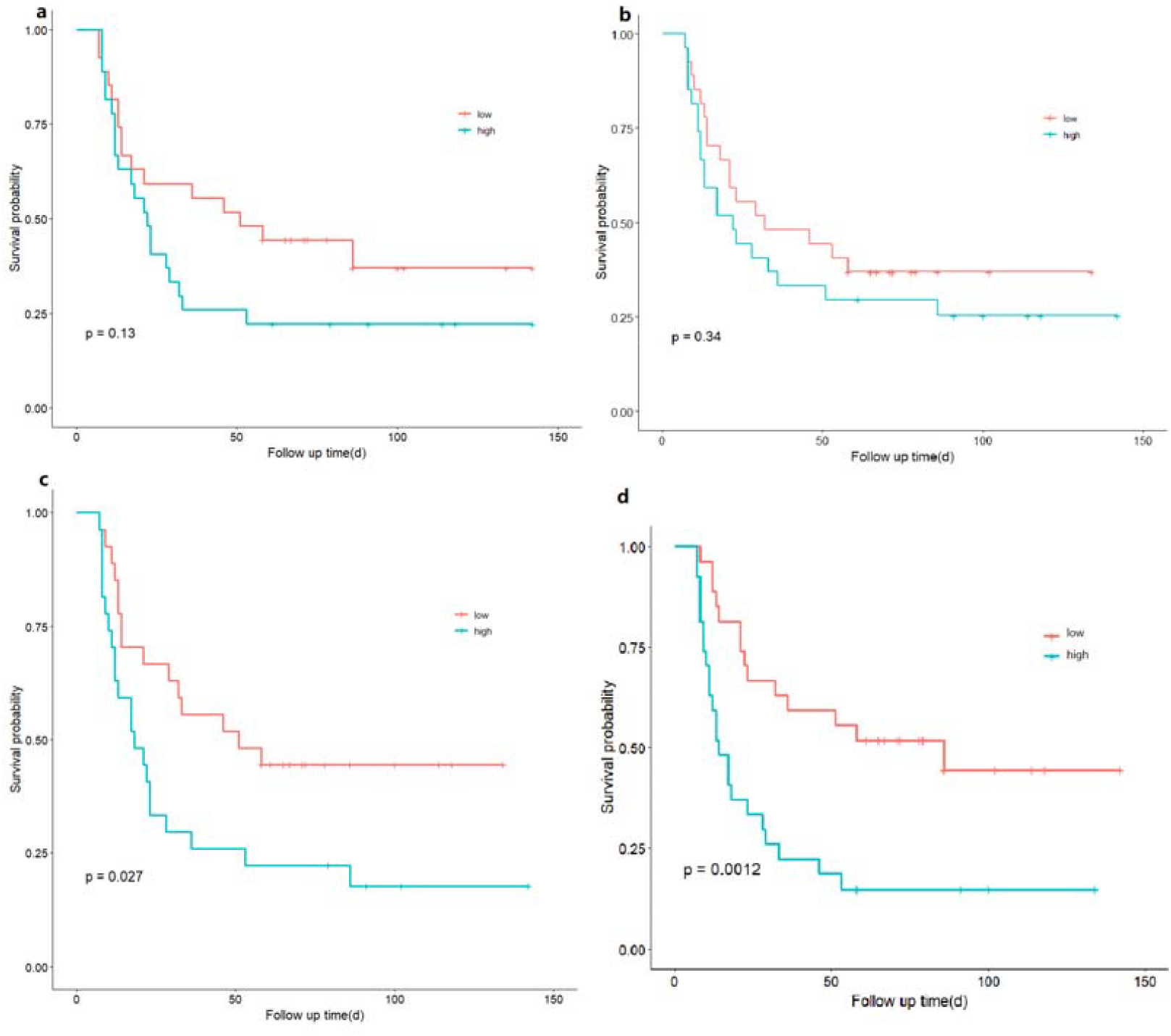
The comparion of KM curves for different models: (a) KM curve for radiomics model; (b) KM curves for baseline CNN model; (c) KM curves for CON-2m model; (d) KM curves for DSN-2m model.

### Interpretation analysis

Figure 4 present the TSNE visualization plot from feature-level (a-b) and patient-level (c-f). Compared with CNN-based deep features (Fig.4a), CON-based deep features (Fig.4b) distribute more densely and show higher accordance with radiomics feature space. While radiomics features distributed in a continuous way, deeply-learned features present different clusters far from each other, indicating that deep network potentially extracts discriminative features from tumor images. Fig.4c showed that the radiomics model split the patients into two groups, while the two groups showed no distinction between DFS time. Different from radiomics model, CNN, CON and DSN model showed significant trends from high DFS to low DFS (Fig.4d-f). However, the patient distributions are very dense in CNN and CON model while the DSN model explicitly separate different patients.

**Figure 4.**
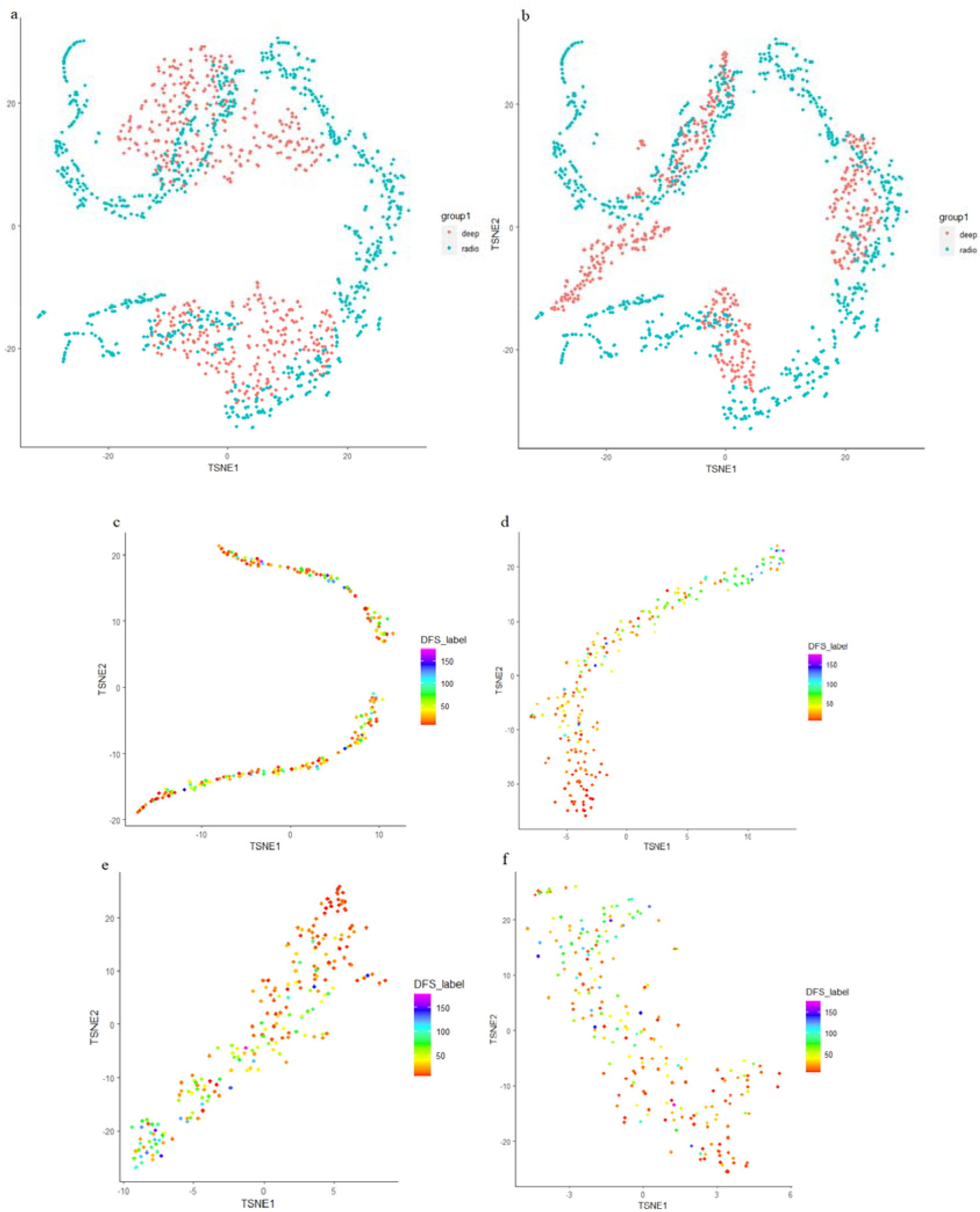
(a) TSNE plot comparing deep features with radiomics features; (b) TSNE plot comparing CON features with radiomics features; (c) TSNE for radiomics model; (d) TSNE for CNN model; (e) TSNE for CON model; (f) TSNE for DSN model

We performed visualization on the last layer of deep networks to reveal what properties have been learned. Grad-CAM visualization on one case with 23 months of DFS showed that, CNN model focused more on the background, while the CON-2m model and DSN-2m model focused more on the tumor lesion (Fig.5). Besides, DSN-2m focused a large part of the tumor while CON-2m focused only a small part of the tumor. This indicated that radiomics-based knowledge guided deep learning to focus more information on tumor lesion, and combining radiomics loss with deep learning loss could further update the model weights to extract more meaningful deep features.

**Figure 5.**
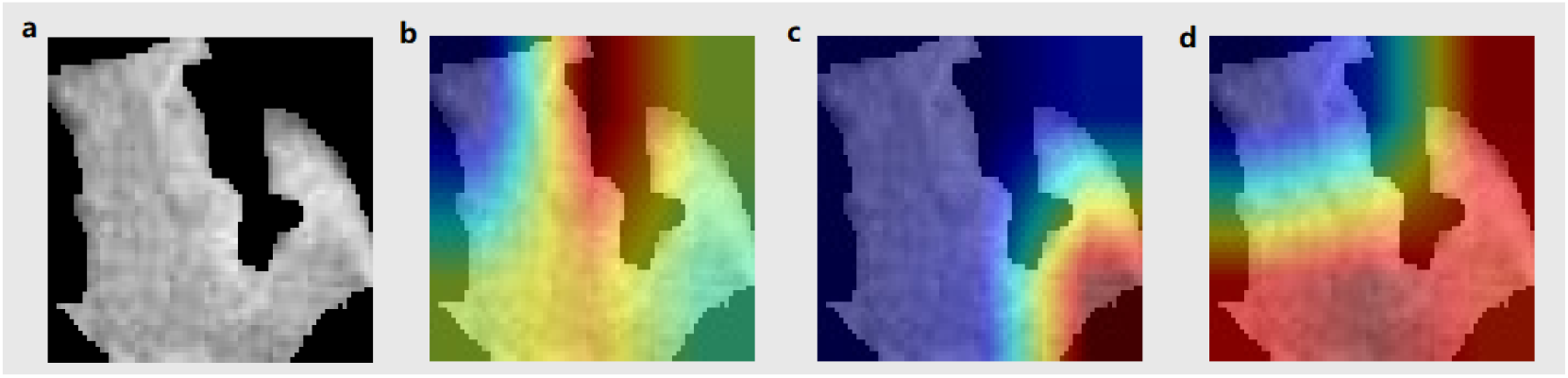
Grad-CAM visualization of the last layer features learned by the deep networks. (a) Grad-CAM plot of baseline CNN; (b) Grad-CAM plot of CON model; (c) Grad-CAM plot of DSN model.

## Discussion

In this study, we developed an integration model which intuitively embedded radiomics features and clinical features into deep learning model to predict the DFS of gastric cancer. We found our proposed model obtained an improvement of C-index of 3%-5%. Interpretation analysis revealed the addition of radiomics features helps deep learning extract useful information to predict disease progression. Our proposed models fully leverage the features extracted from clinical reports and single-phase CT image, which potentially revealed inter-tumor heterogeneity of patients with GC.

Our studies identified new radiomics biomarkers for preoperative DFS prediction. From Supplementary Table S1, tumor size and age are previously reported a biomarker for patient prognosis, and in our clinical model, the features were selected though with no statistical significance. In the model combining radiomics features with clinical features, only one clinical variable (age) was selected. Nine radiomics features were selected in radiomics model, whereas the “ratio” feature was involved. Though the “ratio” feature was roughly calculated by dividing the two-dimension image size, the feature potentially represents the morphology and structure of the tumor. As reported in previous studies, the morphology of cancer is related to the tumor growth patterns and is related to patient prognosis [27, 28]. Our study preliminarily implies the morphology indicated by CT image would be an important factor to DFS prediction of GC patients. However, when integrating deep features with radiomics features, the ‘ratio’ feature has not been selected. This may be due to the rough definition of the morphology. Further investigation is required in the future works.

The deep learning-based integration framework proposed presented several advantages. Firstly, the model can improve the feature extraction ability of deep learning and alleviate the few-shot learning problem. The Grad-CAM visualization plot and T-SNE plot at feature level indicated that the knowledge-guided deep learning model improved the feature extraction ability. Secondly, the model can potentially identify tumor heterogeneity and infer disease progression trajectory. T-SNE plot at sample level showed that the capability of deep learning-based model in identifying disease progression trajectory. While the DSN model also presented disease progression trajectory, the deep features could also extract the difference between different GCs, showing its potentials in identifying inter-tumor heterogeneity. Thirdly, the model is flexible and can integrate different knowledge. The proposed model is an extendable and flexible model which could be fitted to many different scenarios.

Some previous studies have developed and applied similar methods in medical image analysis, including Wei et al [16]. The model proposed by Wei et al is based on variational autoencoder. Variational autoencoder (VAE) [29] could learn the distribution of data, and has been widely applied in many scenarios in recent years including multi-modal learning. However, the data distribution would not be properly learned by the basic VAE in few-shot scenarios, and would induce variations and bias during the model training. Compared to the work, we proposed a CNN-based framework to extract deep features and used fully connected layers to learn the association between radiomics/clinical features with the outcome target, which maximally remains original information of radiomics/clinical features. Moreover, our study provided independent validation of the model, which was not involved in the study mentioned above.

Deep learning possesses the capability of automatic feature extraction, while lacking of interpretability. When dealing with biomedical scenarios, the biology or pathology mechanism revealed behind the model is especially important. In recent years, there is a growing trend of developing biological interpretable models. For image-based modelling, Jiang et al. [30] has established a multi-task deep learning model which predict both the DFS and tumor microenvironment (TME) information for GC. Through the guidance of TME knowledge, the model learned interpretable image features revealing potential biological mechanism for GC prognosis. In our study, the proposed model intuitively incorporate expert-designed radiomics and clinical knowledge into deep network, which would also suggest a future direction on exploring the biological interpretability of deep learning-based medical image analysis.

Beyond the discoveries made, our study has some limitations. Firstly, the samples included were limited, which greatly affect the generalization of deep learning model. Secondly, only tumor region was delineated for each image, while the peritumoral region was also important as it presents the tumor microenvironment [5]. Finally, although we performed interpretation on the meaning of the deep features learned by proposed model, we did not consider more in-depth investigation on how these deep features could be related to biological processes. Our future works will focus on interpreting these imaging-based deep features using paired RNA-seq data with images.

Overall, our study suggests that the integration of clinical parameters, deep learning features, and radiomics features in a deep learning framework could guide deep learning to learn biologically meaningful features and extract the high-level tumor heterogeneity, which finally promote the DFS prediction of gastric cancer.

## Supporting information

From Supplementary Table S1,

## Notes

### Competing Interest Statement

The authors have declared no competing interest.

### Summary of Updates

1.The author list 2.The Discussion part 3.Some other details.

## References

1. Thrift, A.P., T.N. Wenker, and H.B. El-Serag, Global burden of gastric cancer: epidemiological trends, risk factors, screening and prevention. Nature Reviews Clinical Oncology, 2023. 20(5): p. 338–349.

2. Machlowska, J., et al., Gastric Cancer: Epidemiology, Risk Factors, Classification, Genomic Characteristics and Treatment Strategies. Int J Mol Sci, 2020. 21(11).

3. Jiang, Y., et al., Radiomics signature of computed tomography imaging for prediction of survival and chemotherapeutic benefits in gastric cancer. EBioMedicine, 2018. 36: p. 171–182.

4. Wang, S., et al., Preoperative computed tomography-guided disease-free survival prediction in gastric cancer: a multicenter radiomics study. Med Phys, 2020. 47(10): p. 4862–4871.

5. Li, J., et al., Intratumoral and Peritumoral Radiomics of Contrast-Enhanced CT for Prediction of Disease-Free Survival and Chemotherapy Response in Stage II/III Gastric Cancer. Front Oncol, 2020. 10: p. 552270.

6. Hu, C., et al., Deep learning radio-clinical signatures for predicting neoadjuvant chemotherapy response and prognosis from pretreatment CT images of locally advanced gastric cancer patients. Int J Surg, 2023. 109(7): p. 1980–1992.

7. Jiang, Y., et al., Non-invasive tumor microenvironment evaluation and treatment response prediction in gastric cancer using deep learning radiomics. Cell Rep Med, 2023. 4(8): p. 101146.

8. Jiang, Y., et al., Predicting peritoneal recurrence and disease-free survival from CT images in gastric cancer with multitask deep learning: a retrospective study. The Lancet Digital Health, 2022. 4(5): p. e340–e350.

9. Tian, Y., et al., Assessing PD-L1 Expression Level via Preoperative MRI in HCC Based on Integrating Deep Learning and Radiomics Features. Diagnostics (Basel), 2021. 11(10).

10. Bizzego, A., et al. Integrating deep and radiomics features in cancer bioimaging. in 2019 IEEE Conference on Computational Intelligence in Bioinformatics and Computational Biology (CIBCB). 2019.

11. Meng, M., et al. Radiomics-enhanced Deep Multi-task Learning for Outcome Prediction in Head and Neck Cancer. in HECKTOR@MICCAI. 2022.

12. Chen, X., et al., A Combined Model Integrating Radiomics and Deep Learning Based on Contrast-Enhanced CT for Preoperative Staging of Laryngeal Carcinoma. Academic Radiology, 2023. 30(12): p. 3022–3031.

13. Lee, C.-Y., et al., Deeply-Supervised Nets, in Proceedings of the Eighteenth International Conference on Artificial Intelligence and Statistics, L. Guy and S.V.N. Vishwanathan, Editors. 2015, PMLR: Proceedings of Machine Learning Research. p. 562-570.

14. Han, Y., et al., Using Radiomics as Prior Knowledge for Thorax Disease Classification and Localization in Chest X-rays. AMIA Annu Symp Proc, 2021. 2021: p. 546–555.

15. Nanni, L., S. Ghidoni, and S. Brahnam, Handcrafted vs. non-handcrafted features for computer vision classification. Pattern Recognition, 2017. 71: p. 158–172.

16. Wei, L., et al., A deep survival interpretable radiomics model of hepatocellular carcinoma patients. Physica Medica, 2021. 82: p. 295–305.

17. Wang, G., et al., Prediction of Microvascular Invasion of Hepatocellular Carcinoma Based on Preoperative Diffusion-Weighted MR Using Deep Learning. Acad Radiol, 2021. 28 Suppl 1: p. S118–s127.

18. Meng, L., et al., 2D and 3D CT Radiomic Features Performance Comparison in Characterization of Gastric Cancer: A Multi-Center Study. IEEE J Biomed Health Inform, 2021. 25(3): p. 755–763.

19. Abbasian Ardakani, A., et al., Interpretation of radiomics features–A pictorial review. Computer Methods and Programs in Biomedicine, 2022. 215: p. 106609.

20. van Griethuysen, J.J.M., et al., Computational Radiomics System to Decode the Radiographic Phenotype. Cancer Res, 2017. 77(21): p. e104–e107.

21. Simon, N., et al., Regularization Paths for Cox’s Proportional Hazards Model via Coordinate Descent. Journal of Statistical Software, 2011. 39(5): p. 1–13.

22. Katzman, J.L., et al., DeepSurv: personalized treatment recommender system using a Cox proportional hazards deep neural network. BMC Med Res Methodol, 2018. 18(1): p. 24.

23. Yasaka, K., et al., Deep learning with convolutional neural network in radiology. Jpn J Radiol, 2018. 36(4): p. 257–272.

24. Maaten, L.V.D., Accelerating t-SNE using tree-based algorithms. J. Mach. Learn. Res., 2014. 15(1): p. 3221–3245.

25. Maaten, L.v.d. and G.E. Hinton, Visualizing Data using t-SNE. Journal of Machine Learning Research, 2008. 9: p. 2579–2605.

26. Selvaraju, R.R., et al. Grad-CAM: Visual Explanations from Deep Networks via Gradient-Based Localization. in 2017 IEEE International Conference on Computer Vision (ICCV). 2017.

27. Lee, J., et al., Interfacial geometry dictates cancer cell tumorigenicity. Nature Materials, 2016. 15(8): p. 856–862.

28. Almagro, J., et al., Tissue architecture in tumor initiation and progression. Trends Cancer, 2022. 8(6): p. 494–505.

29. Kingma, D.P. and M. Welling, Auto-Encoding Variational Bayes. CoRR, 2013. abs/1312.6114.

30. Jiang, Y., et al., Biology-guided deep learning predicts prognosis and cancer immunotherapy response. Nat Commun, 2023. 14(1): p. 5135.

