## Supplementary material for "An interpretable integration model improving disease-free survival prediction for gastric cancer based on CT images and clinical parameters": From Supplementary Table S1,

**Supplementary Table S1 Features selected by lasso cox regression**

| Model | Features |
| --- | --- |
| Radiomics | "ratio"  "log.sigma.1.0.mm.3D_firstorder_Maximum"  "log.sigma.3.0.mm.3D_firstorder_Maximum"  "log.sigma.3.0.mm.3D_glcm_Idmn"  "log.sigma.3.0.mm.3D_glszm_SizeZoneNonUniformityNormalized"  "log.sigma.4.0.mm.3D_glszm_SmallAreaLowGrayLevelEmphasis"  "wavelet.HH_glrlm_ShortRunHighGrayLevelEmphasis"  "wavelet.LL_glrlm_LongRunLowGrayLevelEmphasis"  "wavelet.LL_ngtdm_Complexity" |
| Clinical | "size1" "CA125" "CEA" "age" |
| Radiomics+Clinical | "ratio"  "log.sigma.1.0.mm.3D_firstorder_Maximum"  "log.sigma.3.0.mm.3D_firstorder_Maximum"  "log.sigma.3.0.mm.3D_glcm_Idmn"  "log.sigma.3.0.mm.3D_glszm_SizeZoneNonUniformityNormalized"  "log.sigma.3.0.mm.3D_glszm_SmallAreaHighGrayLevelEmphasis"  "log.sigma.4.0.mm.3D_glszm_SmallAreaLowGrayLevelEmphasis"  "wavelet.LL_glrlm_LongRunLowGrayLevelEmphasis"  "wavelet.LL_gldm_LargeDependenceLowGrayLevelEmphasis"  "wavelet.LL_ngtdm_Complexity"  "age" |
| Deep | "40" "75" "127" "139" "150" "165" "172" "185" "218" "240" "290" "301" "304" "331" "370" "397" "399" "467" "491" |
| Radiomics+Deep | "log.sigma.1.0.mm.3D_firstorder_Maximum" "wavelet.LH_glcm_Imc1" "wavelet.LL_firstorder_Minimum" "wavelet.LL_glrlm_LongRunLowGrayLevelEmphasis" "wavelet.LL_gldm_LargeDependenceLowGrayLevelEmphasis" "5" "22" "40" "75" "127" "150" "165" "172" "185" "218" "240" "290" "301" "304" "331" "370" "397" "399" "467" "491" |
| Radiomics+Deep+Clinical | "log.sigma.1.0.mm.3D_firstorder_Maximum" "wavelet.LL_firstorder_Minimum" "wavelet.LL_glrlm_LongRunLowGrayLevelEmphasis" "wavelet.LL_gldm_LargeDependenceLowGrayLevelEmphasis" "5" "22" "40" "75" "127" "150" "165" "172" "185" "218" "240" "290" "301" "304" "331" "370" "373" "397" "399" "491" |
